# Description of novel marine bioflocculant-producing bacteria isolated from biofloc of Pacific whiteleg shrimp, *Litopenaeus vannamei* culture ponds

**DOI:** 10.1101/402065

**Authors:** Nurul Fakriah Che Hashim, Nurarina Ayuni Ghazali, Nakisah Mat Amin, Noraznawati Ismail, Nor Azman Kasan

## Abstract

Description of marine bioflocculant-producing bacteria isolated from biofloc of Pacific whiteleg shrimp, *Litopenaeus vannamei* culture ponds was prompted to explore the bacteria that enhanced bioflocculation process in aquaculture wastewater treatment. Certain marine bacteria were potentially secreted extracellular polymeric substances (EPS) which response to the physiological stress encountered in the natural environment that can act as bioflocculants. This study aimed to identify marine bioflocculant-producing bacteria isolated from biofloc; to evaluate their flocculating activities; and to characterize their protein in EPS. Phenotypic and genotypic identification of the bacteria including morphological and molecular approaches were employed, while their flocculating activities were examined via Kaolin clay suspension method and statistically analyzed. The EPS that acted as bioflocculants were extracted using cold ethanol precipitation method. Protein concentration was determined by Bradford assay and protein profiling was finally completed with Sodium Dodecyl Sulfate Polyacrylamide Gel Electrophoresis (SDS-PAGE) method. Six species of marine bacteria known as *Halomonas venusta, Bacillus cereus, Bacillus subtilis, Bacillus pumilus, Nitratireductor aquimarinus* and *Pseudoalteromonas* sp. were successfully identified as bioflocculant-producing bacteria. The highest flocculating activity was exhibited by *Bacillus cereus* at 93%, while *Halomonas venusta* showed the lowest record at 59%. All bioflocculant-producing bacteria species showed different protein concentration that ranged between 1.377 μg/mL to 1.455 μg/mL. Several protein bands with different molecular weight that ranged between 16 kDa to 100 kDa were observed. This study revealed that all the identified bacteria species have high potential characteristics to initiate aquaculture wastewater treatment and may play important roles in bioflocculation process.

**Importance:** Six species of marine bacteria isolated from biofloc of Pacific whiteleg shrimp, *Litopenaeus vannamei* culture ponds were identified as bioflocculant-producing bacteria. Among those six species, *Bacillus cereus*, *Bacillus pumilus*, *Nitratireductor aquimarinus* and *Pseudoalteromonas* sp. were highly potential to be used as booster for rapid formation of biofloc due to their high flocculating activities. Protein content in EPS of novel marine biofocculant-producing bacteria has beneficial consequences on degradation process of organic substances, denitrification of wastes and ions elimination from aquaculture wastes.

## 1. Introduction

Aquaculture is a huge industry that involves cultivation of freshwater and seawater organisms under controlled operations. However, application of effective technologies for wastewater treatment remains minimal in intensive aquaculture operations. High composition of uneaten fish feed and feces in river or sea released by aquaculture operation can cause eutrophication problem (Amirkolaie, 2011). Sludge such as debris, fecal materials and uneaten feed that settled in the bottom sediment can interfere with the interactions of organisms at all biodiversity levels (Yang et al., 2012). Therefore, to ensure long-term sustainability of aquaculture industry, environmental impacts must be minimized and alternative ways such as flocculation process need to be applied.

Flocculation offers an alternative method to overcome the problem of aquaculture wastewater effluent. It was reported as cheap, easy and effective technique to remove cell debris, colloids and suspended particles (Zhang et al., 2012). As compared to other conventional system, this method was volume independent to concentrate dead cells (Salehizadeh & Shojaosadati, 2001). It functioned with the help of flocculant that will alter the nature of suspended particulate materials and enable them to form aggregates or small clumps (Newman, 2011). Flocculants can be divided into synthetic and natural (Yu et al., 2009). For wastewater treatment, synthetic flocculants are the best candidates for aquaculture industry. However, problems regarding their safety status to human health require alternatives flocculants that are more environmental friendly and harmless is crucial to be developed.

Alternatively, green technology metabolites known as bioflocculants which produced by microorganisms can acted similar function as synthetic flocculants to flocculate suspended particles, cells and colloidal solids (Zaki et al., 2011). Many microorganisms including algae, bacteria and fungi isolated from sludge and waste were reported to secrete extracellular polymeric substances. They are mainly consisting of high polymeric substances such as functional proteins, exopolysaccharide, polysaccharides, glycoproteins, protein, nucleic acid and cellulose (Kumar et al., 2004; Feng & Xu, 2008). In other industry, bioflocculants are also widely used as alternative treatment to remove inorganic solid suspensions, dye solutions, food and industrial wastewater (Gao et al., 2009). From other previous studies, there were many bacteria have been reported to be involved in biofloc formation. A bacteria producing an extracellular biopolymer was isolated from contaminated medium and identified as *Bacillus licheniformis* (Xiong et al., 2009). *Paenibacillus* sp. CH11, *Bacillus* sp. CH15, *Herbaspirillium* sp. CH7 and *Halomonas* sp. were reported to produce biopolymer and have been evaluated as bioflocculants in the industrial wastewater effluents treatment (Lin et al., 2012). A strain identified as *Vagococcus* sp. which secreted a large amount of biofloc agents was isolated from wastewater samples collected from Little Moon River in Beijing (Gao et al., 2006). Other bacteria that have been reported as bioflocculant-producing bacteria were *Bacillus firmus* (Salehizadeh & Shojaosadati, 2002), *Citrobacter* spp. TKF04 (Fujita et al., 2001), *Corynebacterium glutamicum* (He et al., 2002), *Enterobacter aerogenes* (Lu et al., 2005), *Nannocystis* sp. Nu-2 (Zhang et al. 2002), *Bacillus subtillis, Bacillus licheniformis, Pacilomyces* sp., and *Nocardia amarae* YK (Deng et al., 2005), *Enterobacter agglomerans* SM 38, *Bacillus subtilis* SM 29 and *Bacillus subtilis* WD90 (Rawhia et al., 2014), *Bacillus cereus* B-11 (Mao et al., 2011), *Serratia ficaria* (Gong et al., 2008), *Lactobacillus delbrukii* sp.*bulgaricus* (Gruter et al., 1993) and *Bacillus alvei* NRC-14 (Abdel Aziz. et al., 2011).

Therefore, the ultimate aim of this study was to characterize the potential bioflocculant-producing bacteria involved in biofloc formation, particularly for aquaculture wastewater treatment.

## 2. Methodology

### 2.1 Location of sampling site

Sampling of biofloc was carried out at Integrated Shrimp Aquaculture Park (iSHARP) Sdn. Bhd (Figure 1). It is located at Setiu, Terengganu (5^°^34’18.32’’N, 102^°^48’25.86’’E), about 30 km away from Universiti Malaysia Terengganu (UMT). iSHARP is a fully Integrated Aquaculture Park developed by Blue Archipelago Berhad specialized for Pacific Whiteleg shrimp, *Litopenaeus vannamei* farming in controlled conditions which operated since 2012. This farm is equipped with superintensive design, biosecurity and vis-à-vis location.

**Figure 1:**
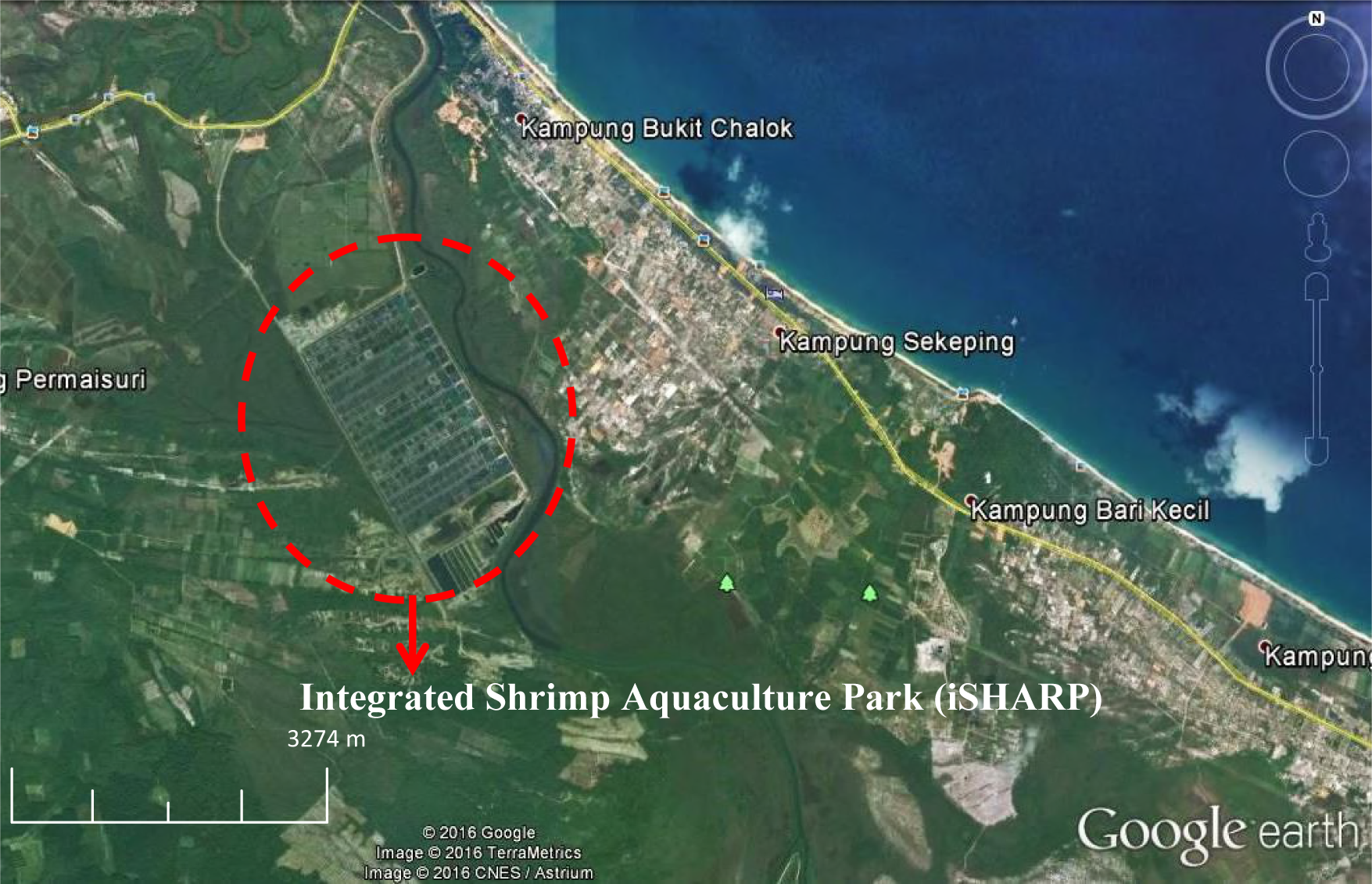
Location of sampling site, Integrated Shrimp Aquaculture Park (iSHARP) Sdn. Bhdin Setiu District, Terengganu, Malaysia (http://www.earth.google.com,2016)

### 2.2 Collection of biofloc samples

Collection of biofloc samples were followed the standard operating procedures (SOP) prepared by iSHARP for biosecurity purpose. Sampling activities were conducted from 25th June 2014 until 29th September 2014. Biofloc samples were collected from fully developed biofloc ponds. In this study, sampling activities were conducted once in every 10 days interval which involved various stage of biofloc formation. For each pond, a total of five litres of pond water containing biofloc samples was collected in pre-acid washed sampling bottles to eliminate contamination and was taken to laboratory for further analysis.

### 2.3 Media preparation

Composition of marine broth contained (per litre): 37.8 g of Difco marine nutrient powder in filtered deionized water. The nutrient agar included (per litre): 55 g of Difco marine agar in filtered deionized water. The Yeast Peptone Glucose (YPG) medium composed (per litre):10.0 g of glucose, 2.0 g of peptone, 0.5 g of urea, 2.0 g of yeast extract, 0.1 g of NaCl, 0.2 g of MgSO_4_.7H_2_O, 0.2 g of KH_2_PO_4_, 5.0 g of K_2_HPO_4_ and 15.0 g of bacteriological agar in filtered deionized water (Ntsaluba et al., 2011). The production medium / enrichment medium (per litre): 10.0 g of glucose, 0.5 g of urea, 0.3 g of MgSO_4_.7H_2_O, 5.0 g of K_2_HPO_4_, 2.0 g of peptone, 0.2 g of KH_2_PO_4_ and 2.0 g of yeast extract in filtered seawater (Cosa et al., 2011). The medium for marine slant agar included (per litre): 10.0 g of glucose, 5.0 g of K_2_HPO_4_, 2.0 g of KH_2_PO_4_, 0.3 g of NH_4_(SO_4_)_2_, 0.5 g of urea, 2.0 g of yeast extract, 0.3 g of MgSO_4_.7H_2_O, 0.1 g of NaCl and 20.0 of agar in filtered deionized water (Gong et al., 2008). All media were adjusted to pH 7 and then sterilized by autoclaving at 121^°^C for 15 min.

### 2.4 Isolation of bioflocculant-producing bacteria from bioflocs

Samples of biofloc were transferred into Imhoff cone for 24 hours to enable the biofloc to settle down. The settled biofloc samples were collected by siphoning out excess water. Biofloc that settled down in Imhoff cone was centrifuged at 6000 rpm for 3 minutes to obtain concentrated biofloc pellet. Concentrated pellet was diluted with saline solution. Isolation of bacteria from biofloc was performed by spread plate method on the surface of marine agar. Biofloc from each pond was plated in 3 replicates. Plates were incubated at 30^°^C for overnight. Single colonies with different morphologies from the cultured plates were inoculated onto new marine nutrient agar plates. The procedure was repeated until pure cultures were obtained.

### 2.5 Screening and identification of bioflocculant-producing bacteria isolated from bioflocs

Screening of bioflocculant-producing bacteria was carried out using production medium and YPG medium. Bioflocculant-producing bacteria were identified through their appearances on solid medium (YPG medium) and liquid medium (production medium). Visual characterization based on ropy, mucoid and slimy was used for identification purposes. Ropy colonies form long filaments when extended with an inoculation loop while mucoid colonies have a glistening and slimy appearance on agar plate (Ortega-Morales et al., 2007). A loop of pure culture of each isolate from marine nutrient agar plate with different colony morphologies were inoculated into 50 mL of marine broth and incubated overnight at 30^°^C for mass production. After incubation, 1 mL of the culture was inoculated into production medium and 0.1 mL was spread evenly on YPG medium. After incubation at 30^°^C for 48 hours, the isolates with ropy morphologies in production medium and mucoid colony morphologies in YPG medium were selected. The isolates were maintained on marine slant agar and kept refrigerated at 4^°^C for further analysis.

### 2.6 Morphological observation and phenotypic characterization of bioflocculant-producing bacteria

Morphological characteristics of bioflocculant-producing bacteria were performed by microscopic observations using Gram staining method. Phenotypic identification was fully carried out according to Bergey’s Manual of Systematic Bacteriology to determine the taxonomy of isolated bioflocculant-producing bacteria as it provided descriptions and photographs of species and tests to distinguish among genera and species (Black, 2005).

### 2.7 Genotypic identification of bioflocculant-producing bacteria through 16S rDNA sequencing

All identified bioflocculant-producing bacteria were further confirmed by genotypic identification through 16S rDNA sequencing.

#### 2.7.1 DNA extraction of bioflocculant-producing bacteria

Identification of microorganisms isolated from biofloc was carried out through molecular approaches. Qiagen DNeasy Blood and Tissue Kit was used to extract bacterial DNA. It was conducted as per manufacturer’s protocol.

#### 2.7.2. DNA quantification and qualification

DNA was quantified using BioDrop μLITE (Isogen, Netherlands). All samples were measured in triplicates and the A_260_/A_280_ ratio values were recorded. Quality of extracted DNA was checked through gel electrophoresis. Gel electrophoresis was conducted according to Mohamad (2014).

#### 2.7.3. Polymerase chain reaction (PCR) amplification

In this study, PCR involved a single set of primer that targets a specific gene that was used to detect an organism. Extracted genomic DNA from individual isolated bacterial strains was subjected to PCR amplification of 16S rDNA using universal PCR primers, 27F and 1492R (Yu et al., 2013) to amplify the 16S rDNA gene. The sequences of primers used were; 27 Forward “5’-AGA GTT TGA TCC TGG CTC AG-3’ “ and 1492 Reverse “5’-ACG GCT ACC TTG TTA CGA CTT-3’ “. PCR was carried out using commercial kit, GoTaq^®^ PCR Core Systems (Promega, USA) for all DNA samples. All PCR reagents used for amplification of bacteria followed recommended reaction volumes and final concentrations provided by manufacturer. Each PCR mixture contained 0.25 μL of Taq polymerase, 10 μL of 10x PCR buffer, 3 μL of MgCl2, 1.5 μL of 200 nM of each primer, 1 μL of 200 nM of dNTP mix, 29.75 μL of distilled deionized water and 3 μL of DNA template (Qiagen, Hilden, Germany). Reactions was carried out in an Eppendorf Mastercycle gradient starting with a denaturation step for 5 minutes at 94^°^C, followed by 35 cycles with 1 cycle consisting of denaturation (94^°^C for 1 minute), annealing (55^°^C for 1 minute), elongation (72^°^C for 2 minutes) and a final extension step for 7 minutes at 72^°^C (Lane, 1991). All PCR products were verified by agarose gel electrophoresis and visualized in gel documentation chamber. Only DNA samples with a single band and clear PCR product shown on agarose gel were selected to be purified and sequenced.

#### 2.7.4 DNA purifications and sequencing

Purification of PCR products was carried out using QIAquick PCR Purification Kit (Qiagen, 28104). The protocol followed manufacturer’s instruction. The amplified PCR products were sent to 1st Base Laboratory, Selangor-Malaysia for sequencing. Obtained 16S rDNA gene sequences were BLAST-analyzed at National Center for Biotechnology Information (NCBI); http://www.ncbi.nlm.nih.gov/BLAST/for similarity search.

### 2.8 Determination of flocculating activity of bioflocculant-producing marine bacteria

All identified bioflocculant-producing marine bacteria were cultured in enrichment medium (Cosa et al., 2011). Inoculum was prepared by incubated in SI-600 Lab Companion Incubator Shaker, with 250 rpm at 30^°^C for 3 days. The resultant culture broth was centrifuged using Hettich Zentrifugen Universal 320 at 8, 000 rpm for 30 minutes at 4^°^C. The cell-free supernatants were used as produced bioflocculant to determine the flocculating activity of the bioflocculant-producing bacteria (Gao et al., 2006).

#### 2.8.1 Flocculating activity of bioflocculant-producing bacteria using Kaolin clay suspension method

Flocculating activity was measured using a modified Kaolin clay suspension method (Kurane et al., 1994). Five gram of kaolin clay was suspended in 1 L of deionized water for preparation of 5.0 g/L of kaolin clay suspension. Kaolin clay suspension was adjusted to pH 7. For flocculating activity, 240 mL of kaolin clay suspension and 10 mL of bioflocculant solution (cell-free supernatant) were added into 250 mL beaker. By using JLT4 Jar/Leaching Tester Velp Scientifica, the flocculating activity was started with rapid mixing at 230 rpm for 2 minutes, followed by slow mixing for 1 minute at a speed of 80 rpm. The stirring speed was reduced to 20 rpm and stirring was continued for 30 minutes. Stirring apparatus was stopped and the samples in the beakers were allowed to settle for 30 minutes. The optical density (OD) of the clarifying solution was measured with Shimadzu UV Spectrophotometer UV-1800 at 550 nm. A control experiment was prepared using the same method but the bioflocculant solution was replaced by deionized water. The experiment was repeated 3 times for each bioflocculant-producing bacteria. The flocculating activity was calculated as follows;

Flocculating activity: [(B-A)/B] × 100% which A and B were the absorbance at 550 nm for sample and reference, respectively.

#### 2.8.2 Statistical analysis

Evaluation on flocculating activity of identified marine bioflocculant-producing bacteria was analyzed using Minitab 16.0 software. One-way ANOVA with grouping information by Tukey Pairwise Comparisons method and 95% confidencelevel was applied. Significant differences between the bacteria were determined at 0.05 level of probability.

### 2.9. Characterization of protein composition in extracellular polymeric substances (EPS) produced by marine bioflocculant-producing bacteria

Each bioflocculant-producing bacteria species was cultured in enrichment medium at 250 rpm in orbital shaker for 3 days at 30^°^C for optimum extracellular polymeric substances (EPS) production (Cosa et al., 2011).

#### 2.9.1 Extraction of EPS from bioflocculant-producing bacteria

A total of 40 mL bioflocculant-producing bacterial culture was treated with 10 mL of 1N NaOH for 30 minutes at 4^°^C before extraction. 1N NaOH treatment was applied to give an effective recovery of EPS and to avoid destruction of EPS. After treatment, culture broth of bacteria was centrifuged at 20,000 rpm for 30 minutes at 4^°^C. After centrifugation process, two layers appeared and the cell-free supernatant layer was taken to extract crude EPS. EPS in the cell-free supernatant fluid was precipitated by addition of 3-volumes of ice cold 95% ethanol. The mixture was later left for 24 hours before it was centrifuged again at 10,000 rpm for 15 minutes (4^°^C). The precipitated EPS was collected on a Whatman filter paper (Grade 1: 11 μm) and precipitated again by addition of 3-volumes of ice cold 95% ethanol and dissolved in water at room temperature for further protein analysis.

#### 2.9.2 Quantification of protein concentration in EPS

Protein in extracted EPS was analyzed for protein concentration by Bradford assay. Bovine Serum Albumin (BSA) was used to prepare a protein standard. Standard containing a range of 1 to 5 μg protein in 100 μL volume were prepared. For blank sample (0 μg/mL), distilled water and dye reagent were used. Each standard solution was pipetted into separate clean test tubes. 5 mL Bradford reagent (Bio-Rad) was added into the standard. The standard then was incubated for five minutes. The absorbance at 595 nm was measured. A standard curve was created by plotting the 595 nm values (y-axis) versus their concentration in μg/mL (x-axis). The same step was repeated for the samples. Finally, the concentration of samples was derived from the standard curve (Bradford, 1976).

#### 2.9.3 Protein profiling by SDS-PAGE

Protein composition in crude EPS was separated by SDS-PAGE. Preparation of sample loading buffer, non-continuous running buffer, isopropanol fixing solution, Coomassie Blue stain solution, resolving gel solution and stacking gel solution for SDS-PAGE were prepared following method described by Laemmli (1970) with a slight of modification. Polyacrylamide gel was cast using 4% stacking gel and 12% resolving gel. The 4% stacking gel was prepared using following reagents; 13.2% v/v of acrylamide/bis solution 37.5:1 (30% T, 2.67% C), 25.2% v/v of stacking buffer (0.5 M Tris-HCl pH 6.8), 1% v/v of Sodium Dodecyl Sulfate (10% w/v), 0.5% v/v of ammonium persulfate (10% w/v), 0.1% v/v of TEMED and the remaining volume was top up with distilled deionized water. The 12% resolving gel was prepared using following reagents: 40% v/v of acrylamide/bis solution 37.5:1 (30% T, 2.67% C), 25% v/v of resolving buffer (1.5 M Tris-HCl pH 8.8), 1% v/v of Sodium Dodecyl Sulfate (10% w/v), 0.5% v/v of ammonium persulfate (10% w/v), 0.5% v/v of TEMED and the remaining volume was top up with distilled deionized water. SDS-PAGE was started with the assembling of glass plate sandwich. Resolving gel solution was poured between the glass plates with a pipette and 1/4 of the space was left free for the stacking gel. The top of the resolving gel was carefully covered with 0.1% SDS solution and left for 30 minutes until the resolving gel polymerized. A clear line has appeared between the resolving gel surface and the solution on top when polymerization was completed. Then the 0.1% SDS solution was discarded and gently washed with double-distilled water. The stacking gel solution was poured carefully with a pipette to avoid formation of bubbles. Combs were inserted and the gel was allowed to polymerize for at least 60 minutes. Combs were removed carefully. The gel was put into the electrophoresis tank. The tank was filled with fresh 1X Tris-glycine-SDS non-continuous running buffer (0.5 M Tris, 1.92 M Glycine, 1% w/v Sodium Dodecyl Sulfate, pH 8.3) to cover the gel wells. Samples were prepared by mixing with sample buffer (0.5 M Tris-HCl, 4% w/v SDS, 20% v/v glycerol, 10% v/v 2-mercaptoethanol, 0.05% w/v bromophenol blue) at ratio 1:1 and were boiled for 10 minutes before loaded into wells. Protein marker, See All Blue Plus (Biorad) was loaded into first lane followed by samples for the rest of lane. Probes were connected and 80 volt power supply was set. The power was increased to 95 volt when the dye reached the resolving gel. SDS-PAGE was stopped when the sign of protein marker reached the foot line of the glass plate. The gel was rinsed with distilled deionized water for two or three times and then isopropanol fixing solution was poured on the gel and let for half an hour. The gel was stained with Coomassie Blue Staining (0.1% w/v Coomassie Brilliant Blue CBR-250, 50% v/v methanol, 10% acetic acid, 40% distilled water) for overnight. After that, the gel was distained with distain solution (10% v/v methanol, 10% v/v acetic Acid, and 80% v/v distilled water) for overnight. At the end, the gel was washed with distilled deionized water with three to four changes over 2-3 hours. The protein band then was viewed using gel documentation system (Biorad).

## 3. Results

### 3.1 Identification of bioflocculant-producing bacteria

In this study, most of the phenotypic characteristics of the isolates were similar to those indicated by Bergey’s Manual of Systematic Bacteriology (Boone et al., 2005). Based on biochemical characterization, the investigated isolates resembled two bacterial genera known as *Bacillus* and *Halomonas*. Two unsuccessfully identified genera were labeled as Unknown sp. 1 and Unknown sp. 2 (Table 1). Six different species that have been identified phenotypically were selected for further genotypic identification through 16S rDNA sequencing. Table 2 showed the purity of the extracted genome from the bioflocculant-producing bacteria prior amplification of the DNA by PCR. The optimum purity ratio of extracted DNA was between 1.7 and 2.0 to ensure that no or less contamination occurred during the extraction process. All isolated bioflocculant-producing bacteria showed an acceptable range of DNA purity and were used as templates in PCR amplification (Figure 2). According to the sequences evaluated in the public databases using BLAST search program on (NCBI) website (http://www.ncbi.nlm.nih.gov/), six species were identified as bioflocculant-producing bacteria from the composition of bacteria isolated from bioflocs. *Halomonas* sp. closely related to *Halomonas venusta*. *Bacillus* sp. 1, *Bacillus* sp. 2 and *Bacillus* sp. 3 closely related to *Bacillus cereus, Bacillus subtilis* and *Bacillus pumilus*, respectively. Unknown sp. 1 closely related to *Nitratireductor aquimarinus* while Unknown sp. 2 closely related to *Pseudoalteromonas* sp. (Table 3).

**Table 1:** Phenotypic characterization of bioflocculant-producing bacteria isolated from biofloc

| Predicted genus | Gram staining | Pigmentation | Shape | Endospore staining | Catalase | Oxidase | Glucose fermentation | Mannitol fermentation | Lactose fermentation | Urease | Indole | Motility | Voges -Proskauer | Citrate | Nitrate reduction | Starch hydrolysis | Phenylalanine deaminase |
| --- | --- | --- | --- | --- | --- | --- | --- | --- | --- | --- | --- | --- | --- | --- | --- | --- | --- |
| *Halomonas* sp. | - | Yellow | Rod | - | + | + | + | + | - | - | na | + | + | + | + | na | na |
| Unknown sp. 1 | - | White | Rod | - | + | + | + | - | - | + | - | - | na | + | + | na | na |
| Unknown sp. 2 | - | White | Rod | - | + | + | + | + | + | na | - | + | na | na | - | na | + |
| *Bacillus* sp. 1 | + | Peach | Rod | + | + | - | + | + | - | - | - | + | + | - | na | na | na |
| *Bacillus* sp. 2 | + | White | Rod | + | + | - | + | - | - | - | - | + | + | - | + | na | na |
| *Bacillus* sp. 3 | + | White | Rod | + | + | + | + | + | + | - | - | + | + | - | - | - | na |

Keys: + = Positive, - = Negative, na = Not applicable

**Table 2:** A_260_/A_280_ ratio of bioflocculant-producing bacteria Rdna

| Genus / Species | A<sub>260</sub>/A<sub>280</sub> ratio |
| --- | --- |
| *Halomonas* sp. | 1.909 |
| *Bacillus* sp. 1 | 1.856 |
| *Bacillus* sp. 2 | 1.923 |
| *Bacillus* sp. 3 | 1.853 |
| Unknown sp. 1 | 1.939 |
| Unknown sp. 2 | 1.906 |

**Table 3:** Sequencing of the 16S rDNA of bioflocculant-producing bacteria isolated from biofloc according to the public databases on National Centre for Biotechnology Information (NCBI)

| Isolated genus | Closest matching strain in NCBI | Sequence similarity (%) | Accession number |
| --- | --- | --- | --- |
| *Halomonas* sp. | *Halomonas venusta* NBRC101901 | 99 | AB681589.1 |
| *Bacillus* sp. 1 | *Bacillus subtilis* YNA61 | 100 | JQ039972.1 |
| *Bacillus* sp. 2 | *Bacillus cereus* MCCC1A06376 | 100 | KJ812466.1 |
| *Bacillus* sp. 3 | *Bacillus pumilus* SH-B9 | 99 | CP011007.1 |
| Unknown sp. 1 | *Nitratireductor aquimarinus* CL-SC22 | 99 | HQ176466.1 |
| Unknown sp. 2 | *Pseudoalteromonas* sp. QY5 | 100 | KP676699.1 |

**Figure 2:**
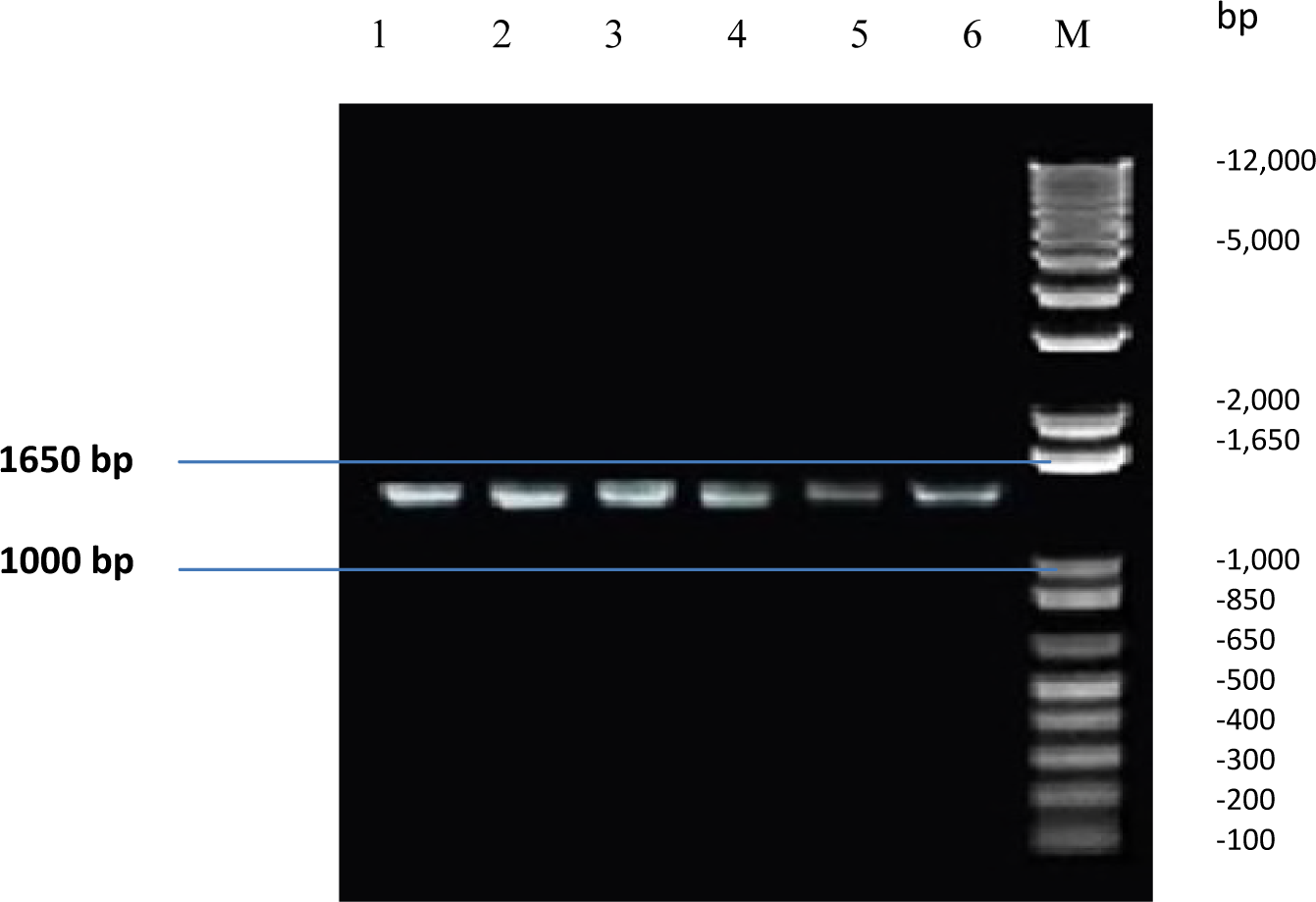
Amplification of ∼1.5 kb fragment of PCR products from bioflocculant-producing bacteria using 1492R and 27F primers. Lane 1: *Halomonas*sp, Lane 2: *Bacillus* sp. 1, Lane 3: *Bacillus* sp. 2, Lane 4: *Bacillus* sp. 3, Lane 5: Unknown sp. 1, Lane 6: Unknown sp. 2 and M: 1kb Plus DNA Ladder (Invitrogen)

### 3.2 The effectiveness of flocculating activity of identified bioflocculant-producing bacteria

Numerically, the highest flocculating activity was showed by *Bacillus cereus* with 93% followed by *Bacillus pumilus* with 92%. *Nitratireductor aquimarinus* showed 89% of flocculating activity and *Pseudoalteromonas* sp. showed 86% of flocculating activity. *Bacillus subtilis* recorded 79% of flocculating activity while *Halomonas venusta* showed lowest record, 59% of flocculating activity. According to statistical analysis using One-Way ANOVA, there was no significant difference (p<0.05) between *Bacillus cereus* (93%) and *Bacillus pumilus* (92%). Besides, there was no significant difference (p<0.05) between *Nitratireductor aquimarinus* (86%) and *Bacillus pumilus* (92%). There was also no significant difference (p<0.05) between *Nitratireductor aquimarinus* (86%) and *Pseudoalteromonas* sp. (86%). According to the statistic, *Bacillus subtilis* was significantly different as well as *Halomonas venusta* (Figure 3).

**Figure 3:**
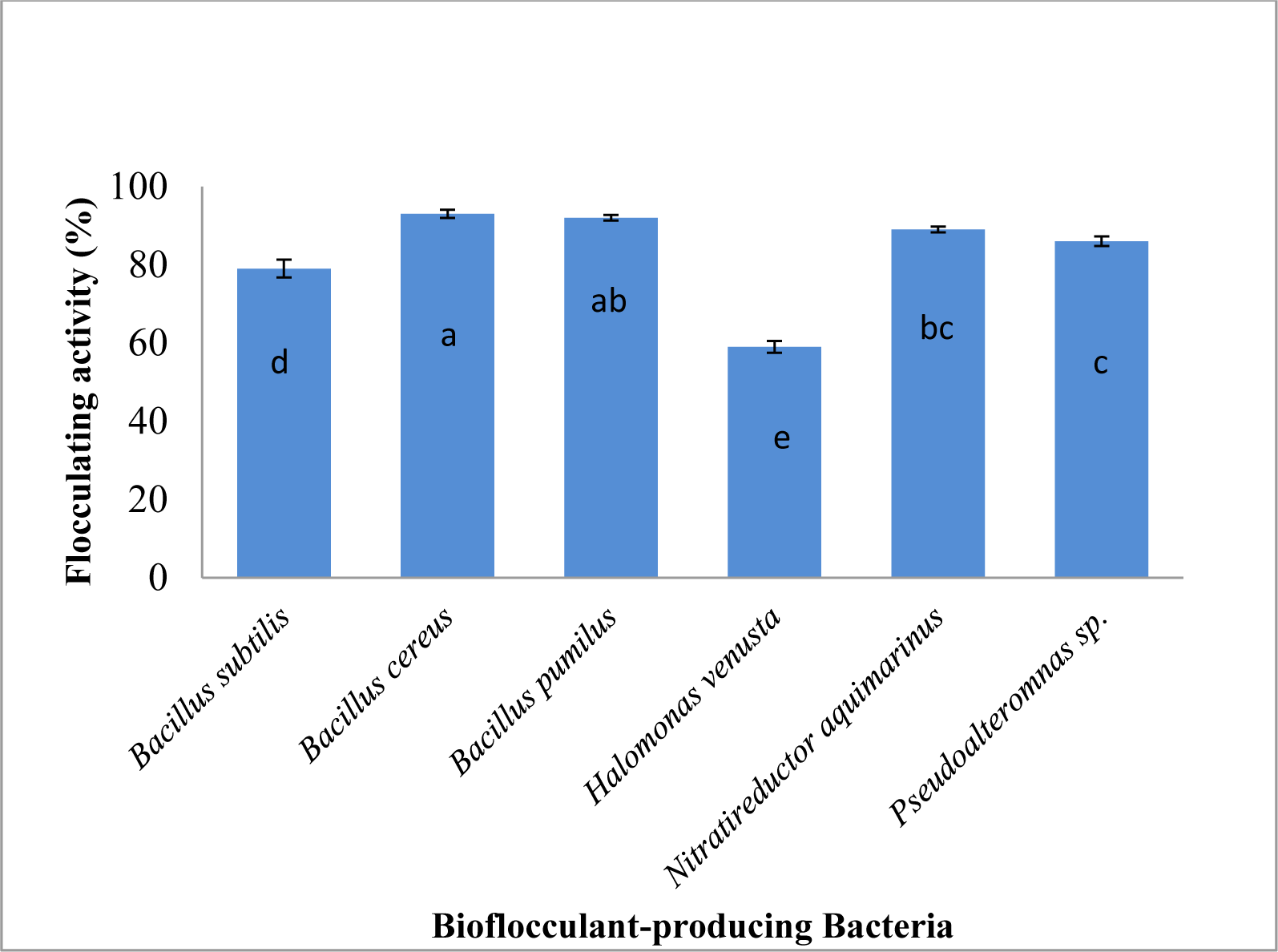
Flocculating activity of bioflocculant-producing bacteria isolated from bioflocs. Note that using grouping information by Tukey Pairwise Comparisons method and 95% confidence, if they do not share the same letter e.g (a, b, c, d, e) it means that they are significantly different. Error bars represented as standard deviation.

### 3.3 Characterization of protein composition in crude extracellular polymeric substances (EPS) from bioflocculant-producing bacteria

Characterization of protein composition in crude EPS from six species of bioflocculant-producing bacteria was analyzed in terms of concentration and molecular weight.

#### 3.3.1 Quantification of protein concentration in crude EPS of bioflocculant-producing bacteria

Each species of bioflocculant-producing bacteria showed different protein concentration (Table 4). The highest protein concentration in extracted EPS was produced by *Bacillus cereus* with 1.455 μg/mL followed by *Bacillus subtilis*, with 1.415 μg/mL. Protein concentration in extracted EPS from *Bacillus pumilus* was 1.403 μg/mL. Protein concentration in extracted EPS from *Pseudoalteromonas* sp., *Halomonas venusta* and *Nitratireductor aquimarinus* were 1.396 μg/mL, 1.388 μg/mL and 1.377 μg/mL respectively.

**Table 4:** Protein concentration in extracellular polymeric substances (EPS) from bioflocculant-producing bacteria

| Biofloculant-producing bacteria | Protein concentration in EPS (µg/mL) |
| --- | --- |
| *B. cereus* | 1.455 |
| *B. subtilis* | 1.415 |
| *B. pumilus* | 1.403 |
| *Pseudoalteromonas* sp. | 1.396 |
| *H. venusta* | 1.388 |
| *N. aquimarinus* | 1.377 |

#### 3.3.2 Protein profiling by SDS-PAGE

Table 5 showed the band of proteins that have been separated by 12% SDS-PAGE at 95V for 1 hour and 30 minutes. Precision PlusProteinTM All Blue Prestained Protein Standard (Biorad) was used as protein marker. Six species of bioflocculant-producing bacteria showed different bands with different molecular weight of protein, ranged between 16 kDa to 100 kDa (Figure 4).

**Table 5:** Protein profiling of marine bioflocculant-producing bacteria on SDS-PAGE

| Lane | Protein marker /<br>Biofloculant-producing bacteria | Estimated molecular weight (kDa) | Number of protein<br>bands |
| --- | --- | --- | --- |
| **A** | *Nitratireductor aquimarinus* | 24-100 | 4 |
| **B** | *Halomonas venusta* | 19-55 | 4 |
| **C** | *Pseudoalteromonas* sp. | 24-55 | 3 |
| **D** | *Bacillus subtilis* | 16-75 | 7 |
| **E** | *Bacillus cereus* | 17-100 | 7 |
| **F** | *Bacillus pumilus* | 18-90 | 6 |

**Lane M** represented Precision PlusProtein™ All blue Prestained Protein Standard (Biorad)

**Figure 4:**
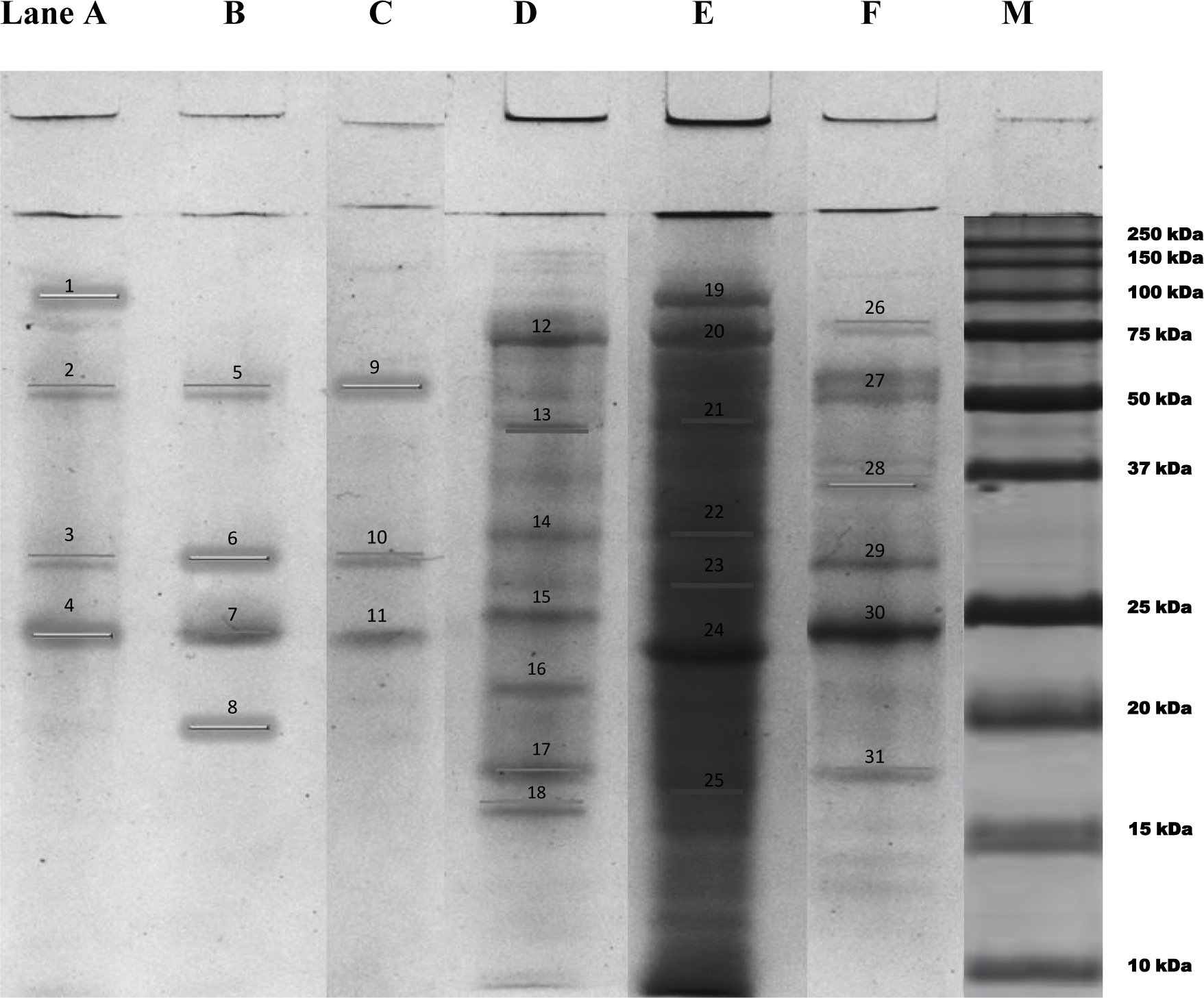
SDS-PAGE profile of extracted EPS from bioflocculant-producing bacteria under denaturing condition on 12% polyacrylamide gel. Lane A: *Nitratireductor aquimarinus*, Lane B: *Halomonas venusta*, Lane C: *Pseudoalteromonas* sp., Lane D: *Bacillus subtilis*, Lane E: *Bacillus cereus*, Lane F: *Bacillus pumilus*, M: Precision PlusProtein^TM^ All Blue Prestained Protein Standard.

## 4. Discussion

### 4.1 Identification of bioflocculant-producing bacteria isolated from bioflocs

From microscopic observation, three identified species were Gram-negative and three species were Gram-positive. The outer membrane of Gram-negative bacteria consists of lipopolysaccharide (LPS), protein and lipoprotein while cell wall of Gram-positive bacteria consists of a thick peptidoglycan layer (Nasir, 2014). The complexity of Gram-negative bacteria cell wall has resulted in better adaptation and survival in marine environment and the outer membrane especially LPS assisted in absorbing nutrient under limited nutrient supply conditions (Nasir, 2014). All Gram-positive bacteria isolated in this study showed positive result in endospore staining test. The capacity to form endospore is unique to certain members of low G-C DNA bases of Gram positive bacteria such as from phylum Firmicutes (Traag et al., 2012). In this study, all species were rod-shaped. According to Thi et al., (2012), Gram-negative and rod-shaped bacteria were dominant in marine environment. Rod-shaped was more advantageous than coccus-shaped because it provides a higher surface-to-volume ratio. Therefore, it is more efficient in nutrient uptake in marine environment (Sjostedt et al., 2012).

In this study, biochemical test was used to identify bacteria species based on differences of biochemical activities of bacteria. It can be carried out conventionally or through commercial identification kit such as API system (Moraes et al., 2013). Commercial identification kit has been widely used as it is fast and its software databases mainly contain clinically important bacteria. It is very useful to identify small number of isolates especially for clinical samples. However, commercial identification kit did not perform well as compared to conventional biochemical test on bacteria species identification. Conventional biochemical test was proven to have accuracy rate of more than 96% while commercial identification kit has only 79 to 94% of accuracy rate because of limited number of tests in commercial identification kit that lead to low percentage of accurate identification (Janda & Abbott, 2002). In this study, species that showed positive result in catalase test indicated that they have the ability to degrade hydrogen through production of catalase (Cappucino & Sherman, 2001). All Gram-negative bacteria in this study contain cytochrome oxidase enzyme because they showed positive result in oxidase test. In carbohydrate fermentation test, species that showed positive result were able to ferment that type of carbohydrate as a carbon source. According to Boone et al., (2005), every species of bacteria even in the same genus has different phenotypic characteristics. This explains why different phenotypic characteristics showed by *Bacillus* sp. in mannitol fermentation test where two species showed positive result while one species showed negative result. In urease test, isolates that showed negative results lack of urease enzyme because they were unable to hydrolyse urea to produce ammonia and carbon dioxide. Urease enzyme is produced by many different bacteria and is reported as virulence factor found in various pathogenic bacteria (Konieczna et al., 2012). In motility test, five species showed positive result. Most bacteria use flagella to move and will enable bacteria to detect and pursue nutrients. Motility is closely linked with chemotaxis which is the ability to orientate along certain chemical gradients (Josenhans & Suerbaum, 2002). In indole test, bacteria that lack enzyme tryptophanase unable to split indole from amino acid tryptophan resulted in no indole production. For Voges-Proskauer (VP) test, isolates that showed positive results can generate acetylbutanediol (ABD) from acetoin. In citrate test, isolates with positive results showed that they were able to utilize citrate as carbon source. This ability depends on presence of a citrate permease enzyme that helps in transport of citrate in the cell (Cappucino & Sherman, 2001). In nitrate reduction test, isolates that showed positive results produced nitrate reductase enzyme because they were capable of reducing nitrate (NO_3_^-^) to nitrite (NO_2_^-^). In phenylalanine deaminase test, isolates that showed positive results was able to remove amino group (-NH_2_) from amino acid with the help of phenylalanine deaminase enzyme. They deaminated phenylalanine and converted it to keto acid, phenylpyruvic acid and ammonia. Isolates that gave positive results in starch hydrolysis test were able to produce extracellular enzymes such as α-amylase and oligo-1,6-glucosidase that hydrolyzed starch. Although biochemical test has been useful for bacteria identification, there were several limitations that need to be considered such as poor reproducibility and difficulties for large-scale applications (Moraes et al., 2013). The best way for identification of bacteria is through conventional biochemical test and 16S rDNA sequencing as no single test identification was proven to have 100% accuracy rate (Janda & Abbott, 2002).

From this study, the result showed that *Bacillus* genus was the most common among the isolates. In previous studies, they were many bacteria of this genus that have been reported as bioflocculant-producing bacteria. For example, *Bacillus licheniformis*, isolated from contaminated medium showed the ability to produce extracellular bioflocculant while *Bacillus* spp. A56 and *Bacillus subtillis* were reported to produce proteinaceous bioflocculants (Xiong et al., 2009; Suh et al., 1997; Deng et al., 2005). In other studies of characterization of microbial EPS, *Bacillus* sp. I-471 and *Bacillus subtilis* DYU1 were identified as bioflocculant-producing bacteria (Kumar et al., 2004; Wu & Ye, 2007). In a study of decolourization of acid dyes, *Bacillus subtilis* and *Bacillus cereus* isolated from disposal site of tannery effluent were identified as bioflocculant-producing bacteria (Anuradha et al., 2014). In a study of role of extracellular polymeric substances in Cu(II) adsorption, the result indicated that the presence of bioflocculant in EPS from *Bacillus subtilis* was significantly enhanced Cu(II) adsorption capacity (Fang et. al., 2011). Besides that, a bioflocculant-producing bacteria known as *Bacillus toyonensis* strain AEMREG6 also has been isolated from sediment samples of a marine environment in South Africa (Okaiyeto et al., 2015). Other genus of *Bacillus* identified as bioflocculant-producing bacteria strains were *Bacillus subtilis* WD90, *Bacillus subtilis* SM29 (Rawhia et al., 2014), *Bacillus alvei* NRC-14 (Abdel Aziz et al., 2011), *Bacillus* sp. CH15 (Lin et al., 2012), *Bacillus firmus* (Salehizadeh & Shojaosadati, 2002) and *Bacillus cereus* B-11 (Mao etal., 2011). All these studies proved that genus of *Bacillus* was one of the most common isolated bioflocculant-producing bacteria.

The genus of *Halomonas* bacteria also showed potential characteristic as bioflocculant-producing bacteria. According to Lin et al., (2012), bioflocculants produced by *Halomonas* sp. were preliminarily evaluated as flocculating agents in the treatment of industrial wastewater effluents. Besides, a bioflocculant-producing bacteria isolated from the bottom sediment of Algoa Bay, South Africa showed 99% of similarity to *Halomonas*sp. Au160H based on 16S rRNA gene sequence. The nucleotide sequence was deposited as *Halomonas* sp. Okoh with accession number HQ875722 (Cosa et al., 2011).

In a study of purification and characterization of EPS with antimicrobial properties from marine bacteria, *Pseudoalteromonas* sp. has been isolated from fish epidermal surface and has been identified as bioflocculant-producing bacteria (Mohd Shahir Shamsir et. al., 2012).

In this study, Unknown sp.1 closely related to *Nitratireductor aquimarinus* when genotypic identification was conducted. *Nitratireductor aquimarinus* has been reported as a bioflocculant-producing bacteria isolated from biofloc of shrimp pond (Nor Azman et al., 2017).

### 4.2 The effectiveness of flocculating activities of identified marine bioflocculant-producing bacteria

Generally, there are factors to be considered in determining the difference of flocculating activity of specific species of bioflocculant-producing bacteria. In this study, cultivation for production of EPS of six identified bioflocculant-producing bacteria was performed following technique of Cosa et al., (2011). The difference of flocculating activity of six identified species of bioflocculant-producing bacteria probably depends on the nature of EPS production during the bacteria growth.

In the present study, glucose, urea and peptone in YPG medium were used as the sources of carbon and nitrogen. It has been reported that carbon and nitrogen sources not only highly manipulate the bioflocculant production and bacterial growth but they also found to play significant roles in flocculating activity (Sheng et al., 2006). From a study of bioflocculant production, glucose was reported to be the ideal carbon source for bioflocculant production by bacteria, as it yielded about 87% flocculating activity compared to sucrose, fructose and starch, which yielded about 75%, 66% and 0% flocculating activities respectively (Sheng et al., 2006). Glucose was reported as the best carbon source to enhance the production of bioflocculants by *Halomonas* sp. V3a (He et al., 2009). For nitrogen source, urea showed the optimal manufacture of bioflocculant and higher flocculating activity compared to peptone (Sheng et al., 2006). Urea was preferred nitrogen source for the cultivation of haloalkalophilic *Bacillus* sp. I-450 (Kumar et al., 2004). According to He et al. (2009) peptone were found to be significant factors that affecting bioflocculant production by *Halomonas* sp. V3a. Previous study of partial characterization and biochemical analysis of bioflocculants of *Halomonas* sp. isolated from sediment, the bioflocculant was optimally produced when glucose and urea were used as sources of carbon and nitrogen (Cosa et. al., 2011).

Initial YPG medium pH that was used for cultivation in this study was pH 7. pH tolerance is another important factor which determine the effectiveness of the bioflocculant in different polluted waters that have wide pH range (Wang et al., 2011). The pH may affect product biosynthesis, cell morphology and structure, cell membrane function, ionic state of substrates, solubility of salts and uptake of various nutrients (Fang & Zhong, 2002). At low pH and high pH, similar effects have been observed where the absorption of H^+^ ions tends to deteriorate the bioflocculant-kaolin complex formation process (He et al., 2010). Maximum bioflocculant producing activity of *Bacillus cereus* and *Bacillus thuringiensis* was affected by pH between pH 7 to pH 8 (Rawhia et al., 2014). However, these observations differ from the result of study carried out by Zheng et al. (2008) in which the maximum flocculating activity of *Bacillus* sp. F19, was observed at pH 2 while *Bacillus* sp. PY-90 was found to be actively high at acidic pH range between 3.0 to 5.0 (Yokoi et al., 1995). *Bacillus toyonensis* strain AEMREG6 exhibited above 60% of flocculating activity at medium pH of 5 (Okaiyeto et al., 2015). Bouchtroch et al., (2001) reported optimal pH values for the flocculating activity of *Halomonas maura* was pH 7.2 and pH 7.0. *Halomonas* sp. V3a also attained the highest flocculating activity at pH 7 (He et al., 2010). In a study of partial characterization and biochemical analysis of bioflocculants of *Halomonas* sp., the bioflocculant was optimally produced with flocculating activity of 87% at pH 7.0 (Cosa et. al., 2011). Most of *Bacillus* bacteria performed very well at acidic pH while *Halomonas* bacteria performed optimally at neutral pH.

Other factor is temperature where flocs formation and floc size distribution caused by the hydrophobic interaction occurs reversibly in response to the change in temperature (Sakohara et al., 2000). In this study, the temperature of bacterial culture was set up at 30^°^C for optimum production of bioflocculant. According to Rawhia et al., (2014), maximum bioflocculant producing activity for *Bacillus cereus* and *Bacillus thuringiensis* was affected by temperature ranged between 30^°^C to 40^°^C and during growth period from 72 hours to 96 hours.

Optimum aeration and dissolve oxygen level during bioflocculant production also important for better bioflocculation performance. Aeration could be beneficial to the growth and performance of microbial cells by improving the mass transfer characteristics with respect to substrate, product or by-product and oxygen (Selale, 2007). To achieve the optimum performance of flocculation, during cultivation of six species of bioflocculant-producing bacteria for bioflocculant production, the orbital shaker was set at 250 rpm to ensure there was dissolved oxygen in the bacteria culture. Besides that, the observed flocculating activity might be due to partial enzymatic deprivation of the polymer flocculant in the late phases of cell growth (Choi et al., 1998).

In this study, *Nitratireductor aquimarinus* shows relatively high flocculating activity comparable to *Bacillus pumilus* and *Pseudoalteromonas* sp. (Figure 3). According to Nor Azman et al., (2017), there is information available about the effectiveness of flocculating activity of *Nitratireductor aquimarinus* as bioflocculant-producing bacteria.

### 4.3 Characterization of protein composition in crude extracellular polymeric substances (EPS) from bioflocculant-producing bacteria

Bioflocculants produced by bioflocculant-producing bacteria were in form of crude extracellular polymeric substances (EPS). Determination of protein concentration in crude EPS is very important to prove that EPS were composed of protein. Protein composition in the crude EPS was believed to enhance the mechanism of bioflocculation. EPS was produced by microorganisms for various purposes in reaction to environmental stresses (Bhatia et al., 2013). Most of bioflocculants by microorganisms were formed during their growth phase. For example, bacteria exploit the nutrients in the culture medium to synthesize high molecular-weight polymeric substances under the action of specific enzymes. Quantity and composition of protein in EPS have been shown to vary depending on bacterial strain and environmental stresses such as temperature, pH and ions (Park & Novak, 2007). Quantification of macromolecules within EPS indicated that proteins and carbohydrates are the major constituents with protein level escalating in EPS as growth proceeded from the exponential phase to the stationary phase (Omoike & Chorover, 2004).

Protein band profile on 12% polyacrylamide gel showed that all bioflocculant-producing bacteria species produced a variety of size and structure of protein in EPS. The ability of proteins to move through the gel is depending on their size and structure and relative to the pores of the gel. Large molecules migrate slower than small molecules and this movement created the separation of distinct particles within the gel. In this study, *Bacillus subtilis, Bacillus cereus* and *Bacillus pumilus* showed a quite intense of protein bands on SDS gel. The protein bands that appeared on SDS gel for *Bacillus subtilis, Bacillus cereus* and *Bacillus pumilus* were ranged between 16 - 75 kDa, 17 - 100 kDa and 18 - 90 kDa respectively. Many studies reported that extracted EPS from *Bacillus* sp. usually are used as stabilizers, emulsifiers, binders, gelling agent and film formers. EPS from *Bacillus* genus had been an interesting topic because they are Generally Recognized as Safe (GRAS). Chemical compositions in EPS such as proteins, neutral polysaccharides, amphiphilic molecules and charged polymers that produced by wild-type *Bacillus subtilis* strains cultured under controlled laboratory conditions reveal a wide range of molecular weight with sizes ranging from 0.57 kDa to 128 kDa (Omoike & Chorover, 2004). Most of proteins are found freely in the surrounding medium as they dissociated from cells and some are found within exopolymeric matrix. Proteins that composed by *Bacillus subtilis* also included the proteins that responsible for the extracellular enzymes discharge and protein export from the cytoplasm to the surrounding environment. Many proteins that secreted by *Bacillus subtilis* also involved in the degradation of molecules such as extracellular nucleic acids, phytic acid, lipids and glutathione (Tjalsma et al., 2004). In a study of production and characterization of EPS from bacteria isolated from pharmaceutical laboratory sinks by Nanda & Raghavan, (2007), molecules, proteins and functional groups are found in the EPS produced from *Bacillus subtilis* using FTIR analysis. The biopolymer flocculants named FQ-B1 and FQ-B2, produced by *Bacillus cereus* and *Bacillus thuringiensis* were precipitated by chemical elemental analysis and UV scan were performed for investigating the purified bioflocculant contained 2.56 μg/ mL (83.01%) and 1.78 μg/ mL (84.73%) of protein respectively (Rawhia et al., 2014). In a study of glycoprotein bioflocculant, chemical analysis showed that purified bioflocculant produced by *Bacillus toyonensis* strain AEMREG6 was mainly composed of polysaccharide (77.8%) and protein (11.5%) (Okaiyeto et al., 2015). Extracted bioflocculants from *Bacillus subtillis* can be used as an alternative agent to eliminate copper at lower concentrations but further study needs to be carried out on its actions mechanism, scaling up process and modifications to enhance its ability in order to make it more reliable for industrial utilization (Azmi et al., 2015).

In this study, even though *Halomonas venusta, Pseudoalteromonas* sp. and *Nitratireductor aquimarinus* did not show very high concentration of protein in their extracted EPS, they still showed several prominent protein bands. *Halomonas venusta* showed four prominent protein bands that ranged between 19 - 55 kDa. It showed that protein was one of the main compositions in its bioflocculants. This study was supported by a study of partial characterization of *Halomonas* sp. where chemical analysis revealed that bioflocculant produced by *Halomonas* sp. was mainly polysaccharide and protein (Cosa et. al., 2011).

*Pseudoalteromonas* sp. showed three prominent protein bands that ranged between 24 - 55 kDa. It showed that protein was one of the components in its EPS. Previous finding on purification and characterization of EPS with antimicrobial properties from *Pseudoalteromonas* sp. has revealed that up to eight protein types of unknown proteins were detected within the EPS, with size of molecular weight ranging from 15.486 kDa to 113.058 kDa (Mohd Shahir Shamsir et. al., 2012). The *Pseudoalteromonas* sp. in the study also showed to produce the highest amount of EPS during the first 24 hours of culture.

The result obtained in the present study suggests that *Nitratireductor aquimarinus* is a potential bioflocculant-producing bacteria. These bacteria produce four prominent protein bands that ranged between 24 - 100 kDa when analyzed using SDS-PAGE. However, there was no study of EPS characterization to indicate and support that its proteins from EPS can act as bioflocculant since no study claimed *Nitratireductor aquimarinus* as bioflocculant-producing bacteria.

## 5. Conclusion

Six species of marine bacteria were successfully identified as bioflocculant-producing bacteria from bioflocs. They were closely similar to *Halomonas venusta, Bacillus cereus, Bacillus subtilis, Bacillus pumilus, Nitratireductor aquimarinus* and *Pseudoalteromonas* sp. The group of high flocculating activity was exhibited by *Bacillus cereus* (93%), *Bacillus pumilus* (92%), *Nitratireductor aquimarinus* (89%) and *Pseudoalteromonas* sp. (86%). *Bacillus subtilis* (79%) represented group of intermediate flocculating activity while *Halomonas venusta* (59%) was categorized as group of low flocculating activity. For protein characterization of crude EPS, all species of bioflocculant-producing bacteria have different protein concentration that ranged between 1.377 μg/mL to 1.455 μg/mL with different banding patterns between three to seven bands at different molecular weight that ranged between 16 to 100 kDa.

It is recommended to further characterize on EPS produced by *Nitratireductor aquimarinus* especially in terms of function and structural using latest advanced methods such as nuclear-magnetic resonance (NMR) to characterize polysaccharide composition and high performance liquid chromatography (HPLC) to separate components of mixture from one another. The methods may assist in order to detect other complex compositions reported in EPS such as polysaccharides, nucleic acid, uronic acid, phospholipid and glycoprotein. The results would be an initial step towards the utilization and modification of EPS in future research in the production of valuable properties especially in aquaculture industry.

## 6. Acknowledgements

This project was supported by the Ministry of Education, Malaysia (MOE) under Fundamental Research Grant Scheme, FRGS (vot no. 59401). We also would like to thank iSHARP, Blue Archipelago Berhad, Setiu, Terengganu, Malaysia for *L. vannamei* aquaculture facilities. Finally, to all lab staffs at the Institute of Tropical Aquaculture (AKUATROP), Universiti Malaysia Terengganu who have major contributions throughout the study periods.

